# Mechanotransduction activity facilitates hair cell toxicity caused by the heavy metal cadmium

**DOI:** 10.1101/812438

**Authors:** Caleigh Schmid, Isabella Alampi, Jay Briggs, Kelly Tarcza, Tamara M Stawicki

## Abstract

Hair cells are sensitive to many insults including environmental toxins such as heavy metals. We show here that cadmium can consistently kill hair cells of the zebrafish lateral line. Disrupting hair cell mechanotransduction genetically or pharmacologically significantly reduces the amount of hair cell death seen in response to cadmium, suggesting a role for mechanotransduction in this cell death process, possibly as a means for cadmium uptake into the cells. Likewise, when looking at multiple cilia-associated gene mutants that have previously been shown to be resistant to aminoglycoside-induced hair cell death, resistance to cadmium-induced hair cell death is only seen in those with mechanotransduction defects. In contrast to what was seen with mechanotransduction, significant protection was not consistently seen from other ions previously shown to compete for cadmium uptake into cells or tissue including zinc and copper. These results show that functional mechanotransduction activity is playing a significant role in cadmium-induced hair cell death.

## Introduction

Hearing loss is one of the most common sensory disorders affecting upwards of 20% of Americans over the age of 12 (Goman and Lin, 2016; Lin et al., 2011). One common cause of hearing loss is the death of sensory hair cells. This is also a cause of vestibular dysfunction. Hair cells are sensitive to numerous insults including loud noises, certain therapeutic drugs, and aging (Cheng et al., 2005; Kurabi et al., 2017; Schacht et al., 2012; Yamasoba et al., 2013). There is also evidence that environmental toxins such as heavy metals can lead to hearing loss (Choi and Kim, 2014; Schaal et al., 2017), though this form of hearing loss has not been as well-established or studied as other forms.

Heavy metal toxicity, in general, is a growing environmental concern. Heavy metals are naturally occurring and while some are required in the body at trace levels, higher levels are often associated with toxicity. Other heavy metals, such as cadmium, have no known function or benefit to animals. Cadmium is used in batteries and some pigments and is produced as a by-product of zinc mining. It can be released into the environment through industrial run-off from a variety of different sources including mine drainage water, sewage treatment plans, and hazardous waste sites (IARC, 2012). Working in certain industries comes with a risk of occupational exposure to cadmium, and nonoccupational exposures can occur through diet or smoking (Faroon et al., 2012). Elevated levels of blood or urinary cadmium correlate with a number of health problems including dysfunction of the kidneys, liver, cardiovascular system, osteoporosis, and cancer (Ferraro et al., 2010; Gallagher et al., 2008; García-Esquinas et al., 2014; Hyder et al., 2013; Tellez-Plaza et al., 2012).

Studies looking at the link between cadmium exposure and hearing loss have had conflicting results. Some studies have shown increased hearing thresholds or balance impairments in individuals with higher blood cadmium levels (Choi et al., 2012; Choi and Park, 2017; Min et al., 2012). However, other studies have failed to show a significant association between elevated urinary or blood cadmium levels and hearing loss (Kang et al., 2018; Liu et al., 2018; Shiue, 2013). Some of these conflicting results may come from the different methods of quantifying cadmium levels and hearing loss used by the different groups. Studies in rodents have likewise drawn varying conclusions over whether or not cadmium exposure can cause hearing loss. Hair cell death in organ of Corti cultures and changes in auditory brain response (ABR) and distortion product otoacoustic emissions (DPOAEs) have been seen in mice and rats by some groups (Kim et al., 2008; Liu et al., 2014; Ozcaglar et al., 2001). However, other groups have failed to see vestibular dysfunction or hearing loss following exposure to cadmium alone (Carlson et al., 2018; Klimpel et al., 2017; Whitworth et al., 1999). Again different groups have used different cadmium treatment paradigms which may be responsible for the conflicting results. Defects in auditory and lateral line systems have been more consistently seen in fish following cadmium exposure (Baker and Montgomery, 2001; Faucher et al., 2006, 2008; Low and Higgs, 2015; Montalbano et al., 2018; Sonnack et al., 2015), making fish a useful model to study the mechanisms by which cadmium may cause hair cell death. The lateral line is a superficial sensory structure in aquatic animals that is used to detect water movements and contains hair cells similar to those used for hearing and balance in the mammalian inner ear (Larsson, 2012). It has previously been shown that hair cells of the zebrafish lateral line system are sensitive to many of the same insults as mammalian hair cells (Harris et al., 2003; Ou et al., 2007; Ton and Parng, 2005), and the presence of lateral line hair cells on the surface of the animal facilitates both observation of hair cells and the access of potential ototoxic drugs.

One open question regarding cadmium-induced hair cell death is how cadmium enters hair cells. Cells do not have designated cadmium channels or transporters as cadmium has no function in the cell. Instead, studies looking at other cell types have found that cadmium enters through channels and transporters for other ions. Studies have shown that cadmium can enter cells via zinc transporters ZIP8 and ZIP14, the iron transporter DMT1, the organic cation channels OCT1 and OCT2, and the TRP channels TRPV5, TRPV6, and TRPM7 (Bannon et al., 2003; Dalton et al., 2005; Fujishiro et al., 2009; Girijashanker et al., 2008; Kovacs et al., 2011, 2013; Lévesque et al., 2008; Martineau et al., 2010; Olivi et al., 2001; Soodvilai et al., 2011). It has also been shown that some metals such as copper and zinc can compete with cadmium for uptake into tissues, presumably due to the use of shared uptake routes (Barbier et al., 2004; Komjarova and Bury, 2014). Another potential uptake route for cadmium into hair cells is the hair cell mechanotransduction channel. Other hair cell toxicants including aminoglycoside antibiotics and the chemotherapeutic cisplatin have been shown to require functional mechanotransduction to enter hair cells (Alharazneh et al., 2011; Marcotti et al., 2005; Thomas et al., 2013).

We have found that cadmium can consistently kill hair cells of the zebrafish lateral line system in a dose-dependent manner. This hair cell death is reduced following both genetic and pharmacological inhibition of hair cell mechanotransduction suggesting that mechanotransduction plays a role in cadmium-induced hair cell death, potentially as the route through which cadmium enters hair cells. In contrast to this, we did not see consistent significant protection from cadmium-induced hair cell death when cotreating fish with either zinc or copper, suggesting that these ions are not competing with cadmium for hair cell entry.

## Materials and Methods

### Animals

All experiments used five-day post-fertilization (dpf) *Danio rerio* (zebrafish) larvae. Experiments were carried out with either *AB wild type zebrafish, *cdh23*^*tj264*^ (Nicolson et al., 1998; Söllner et al., 2004), *ift88*^*tz288*^ (Brand et al., 1996; Tsujikawa and Malicki, 2004), or *cc2d2a*^*w38*^ (Owens et al., 2008) mutants. Mutant alleles were maintained in the *AB background and experiments were carried out on offspring of incrosses of heterozygous parents comparing homozygous mutants to both homozygous and heterozygous wild-type siblings from the same clutch. Mutants were separated based on secondary phenotypes; vestibular defects in the case of *cdh23* mutants and body morphology defects in the case of *ift88* and *cc2d2a* mutants.

Larvae were raised in petri dishes containing embryo media (EM) consisting of 1 mM MgSO_4_, 150 μM KH_2_PO_4_, 42 μM Na_2_HPO_4_, 1 mM CaCl_2_, 500 μM KCl, 15 mM NaCl, and 714 μM NaHCO_3._ They were housed in an incubator maintained at 28.5°C. The Lafayette College or University of Washington Institution Animal Care and Use Committee approved all experiments.

### Drug Treatment

Fish were treated with cadmium chloride hemipentahydrate (Fisher Scientific) dissolved in EM for three hours at 28.5°C. Some fish were additionally treated with benzamil hydrochloride hydrate (Sigma-Aldrich), zinc sulfate heptahydrate (Sigma-Aldrich), or copper sulfate pentahydrate (Sigma-Aldrich). For all treatments, fish were put into 6 well plates containing net well inserts and the net wells were moved to plates containing the different solutions the fish were exposed to. Following cadmium treatment, animals were washed three times in EM, euthanized with MS-222 and immediately fixed for immunostaining.

### Immunostaining and Hair Cell Counts

Fish used for immunohistochemistry were fixed for either two hours at room temperature or overnight at 4°C in 4% paraformaldehyde. Antibody labeling was carried out as previously described (Stawicki et al., 2014). Fish used for hair cell counts were labeled with a rabbit anti-parvalbumin primary antibody (ThermoFisher, PA1-933) diluted at 1:1,000 in antibody block (5% goat serum in PBS, 0.2% Triton, 1% DMSO, and 0.2% BSA). Hair cells were counted in the OP1, M2, IO4, O2, MI2, and MI1 neuromasts (Raible and Kruse, 2000) and then an average number of hair cells/neuromast was calculated. Statistics were calculated in GraphPad Prism 6.

## Results

### Acute cadmium treatment can kill hair cells of larval zebrafish in a dose-dependent manner

To investigate potential mechanisms of cadmium uptake into hair cells we first worked to develop an acute cadmium treatment paradigm that would reliably kill hair cells. Cadmium has previously been shown to kill hair cells in both adult and larval zebrafish (Montalbano et al., 2018; Sonnack et al., 2015; Wang and Gallagher, 2013); however, none of these experiments quantified hair cell death in larval fish following acute treatment paradigms. We found that after a three-hour treatment with cadmium, we could see a dose-dependent decrease in lateral line hair cell numbers (Figure 1). This treatment paradigm was thus used in subsequent experiments looking for protection from cadmium-induced hair cell death.

**Figure 1:**
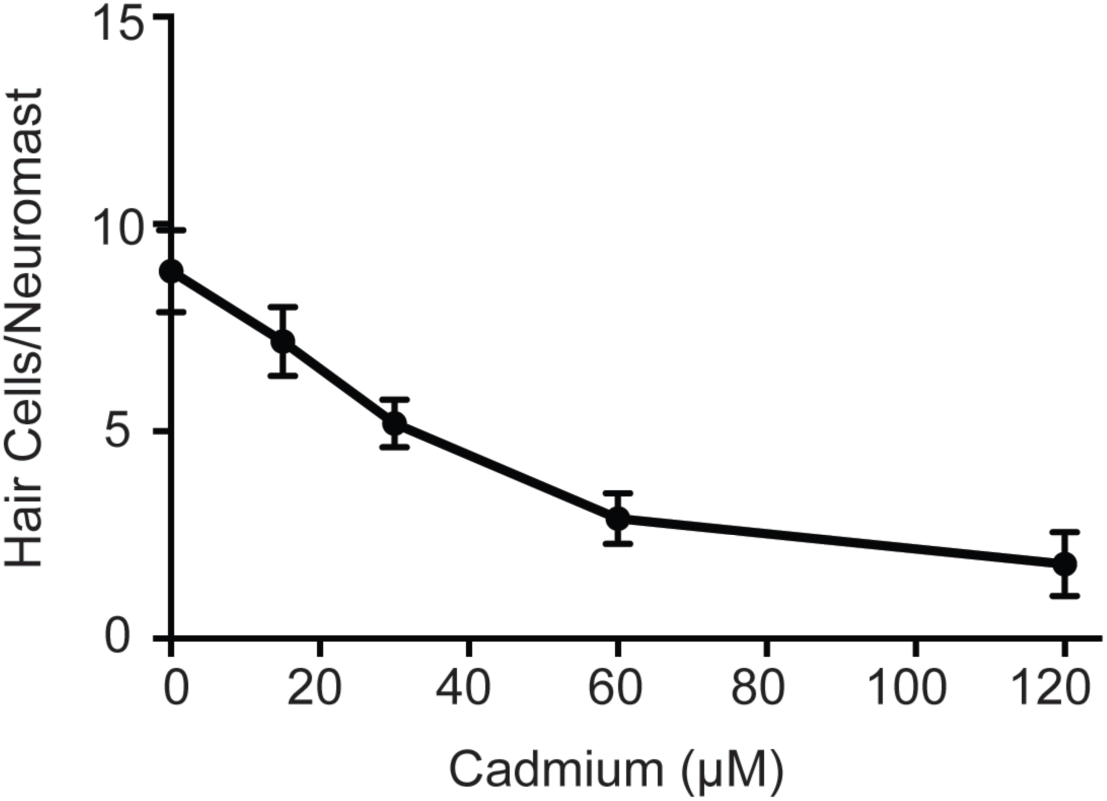
Cadmium kills hair cells in zebrafish larvae in a dose-dependent manner. Treating 5-day post-fertilization (dpf) zebrafish with cadmium doses ranging from 0 - 120 μM for three hours showed an increasing amount of hair cell death with increasing cadmium dose. Data are displayed as mean ± standard deviation. Fish were fixed and stained with parvalbumin and hair cells from six neuromasts were counted (OP1, M2, IO4, O2, MI2, and MI1).

### Impaired mechanotransduction activity reduces hair cell death in response to cadmium

Multiple hair cell toxicants have been shown to enter hair cells in a mechanotransduction-dependent manner (Alharazneh et al., 2011; Marcotti et al., 2005; Thomas et al., 2013) and manipulations that decrease hair cell mechanotransduction activity therefore protect hair cells from these toxicants (Seiler and Nicolson, 1999; Stawicki et al., 2014; Thomas et al., 2013; Wang and Steyger, 2009). To test if impairing mechanotransduction would likewise protect hair cells from cadmium-induced hair cell death, we both genetically and pharmacologically manipulated hair cell mechanotransduction activity in zebrafish larvae. First, we tested *cadherin 23* mutants, also known as *sputnik*. Cadherin 23 is one of the proteins that make up the tip links that link neighboring stereocilia in hair cells. Mutants no longer have tip links and thus do not have functional mechanotransduction activity (Nicolson et al., 1998; Söllner et al., 2004). We found that there was significantly less cadmium-induced hair cell death in these mutants at all doses tested though hair cell death was not eliminated at higher cadmium doses (Figure 2A). We next pharmacologically inhibited mechanotransduction activity by cotreating fish with cadmium and 200 μM benzamil, an analog of amiloride that has been shown to block hair cell mechanotransduction activity (Hailey et al., 2017). We again saw significant protection against cadmium-induced hair cell death at all cadmium doses (Figure 2B).

**Figure 2:**
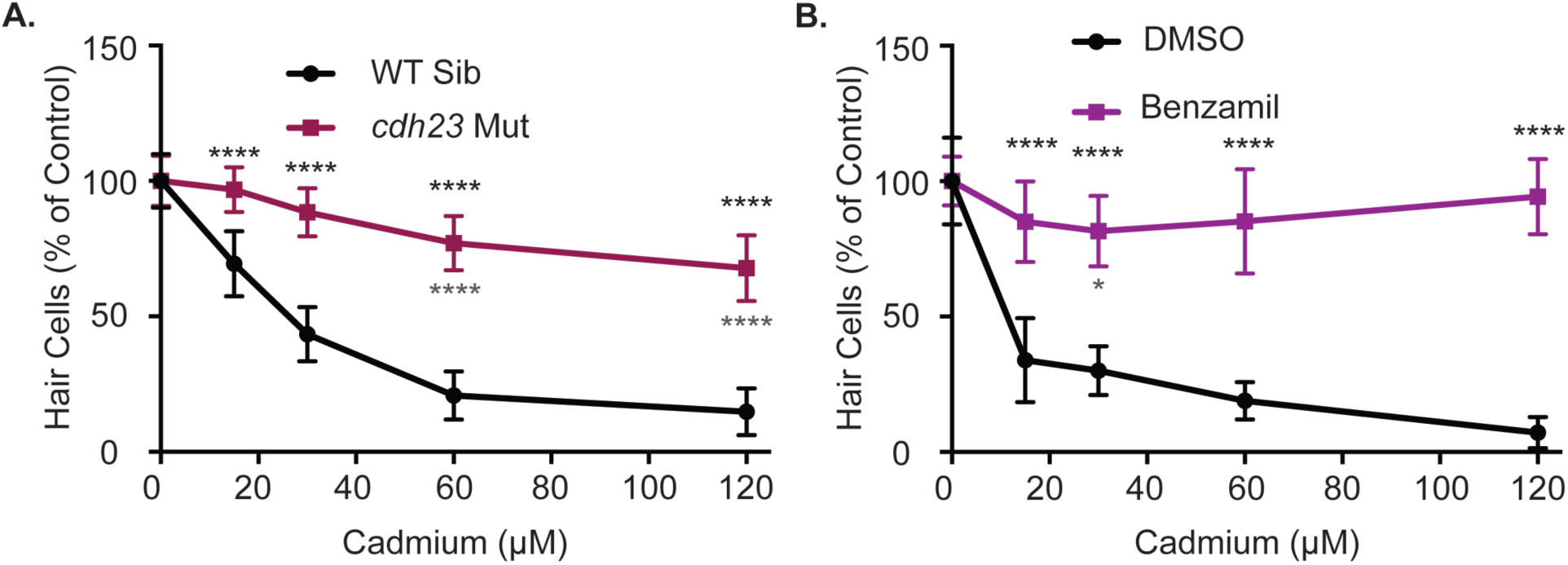
Cadmium toxicity is dependent on functional mechanotransduction activity. (A) *cdh23* mutants, which lack functional mechanotransduction activity, are resistant to cadmium-induced hair cell death. (B) Benzamil, a mechanotransduction blocker, significantly blocks cadmium-induced hair cell death. Data is normalized to the 0 cadmium control for each treatment group. Data are displayed as mean ± standard deviation. * = p<0.05, **** = p<0.0001 by Two-Way ANOVA and Šídák multiple comparisons test. Black stars above the error bars denote significant differences in *cdh23* mutants or benzamil treated fish as compared to controls treated with the same dose of cadmium. Gray stars below the error bars denote significant differences comparing different cadmium doses in *cdh23* mutants or benzamil treated fish to the 0 cadmium control group.

We next tested whether genetic mutants identified through a screen looking for mutants resistant to aminoglycoside-induced hair cell death (Owens et al., 2008) would also be resistant to cadmium-induced hair cell death. We tested two different cilia-associated gene mutants. One, *ift88*, is believed to be resistant to hair cell death in response to the aminoglycoside neomycin due to reduced neomycin uptake as a result of reduced mechanotransduction activity. The other, *cc2d2a*, appears to have normal neomycin uptake and mechanotransduction activity (Stawicki et al., 2016). We found reduced hair cell death in response to cadmium in *ift88* mutants (Figure 3A), further suggesting that mechanotransduction plays a role in cadmium-induced hair cell death. In contrast, we saw no reduction of hair cell death in *cc2d2a* mutants in response to cadmium (Figure 3B).

**Figure 3:**
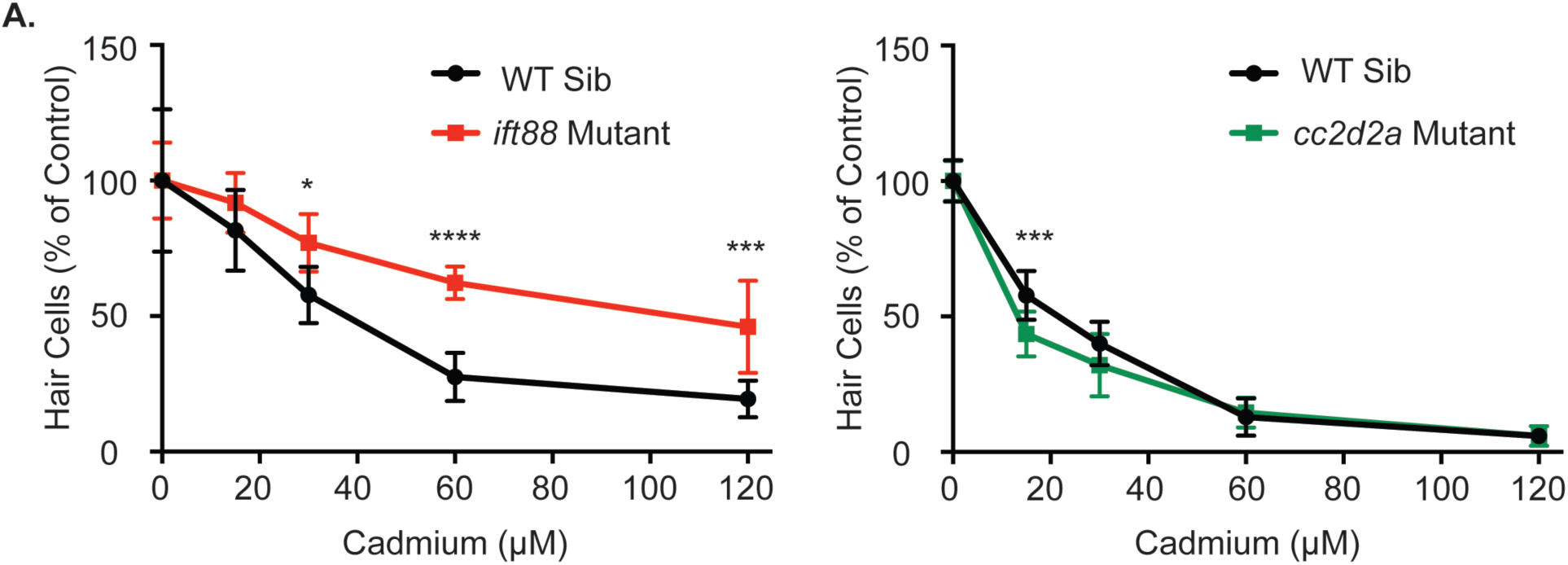
Genetic mutants resistant to neomycin-induced hair cell due to impaired mechanotransduction activity are also resistant to cadmium-induced hair cell death. (A) *ift88* mutants which have previously been shown to have reduced mechanotransduction activity are partially resistant to cadmium-induced hair cell death. (B) *cc2d2a* mutants, which appear to have normal mechanotransduction activity, are not resistant to cadmium-induced hair cell death. Data is normalized to the 0-cadmium control for each treatment group. Data are displayed as mean ± standard deviation. * = p<0.05, *** = p<0.001, **** = p<0.0001 by Two-Way ANOVA and Šídák multiple comparisons test.

### Zinc and copper cotreatment do not protect hair cells from cadmium-induced hair cell death as effectively as impairing mechanotransduction does

As we still saw some hair cell death in *cadherin23* mutants, which completely lack mechanotransduction, we wanted to test if there were alternative means by which cadmium could enter hair cells. Previous experiments have shown that cadmium enters other cell types through zinc transporters (Dalton et al., 2005; Fujishiro et al., 2009), and that zinc can block cadmium uptake (Barbier et al., 2004; Girijashanker et al., 2008). Zinc has also previously been shown to protect against impaired behavioral responses to odorants caused by cadmium in larval zebrafish (Heffern et al., 2018) and to protect against impaired auditory responses caused by cadmium in rats (Agirdir et al., 2002). To test whether zinc could likewise protect against cadmium-induced hair cell death in zebrafish, we co-treated fish with 30 μM of cadmium and varying doses of zinc, based on doses that had previously been shown to protect against cadmium-induced defects in olfactory behavior in larval zebrafish (Heffern et al., 2018), with and without a one-hour zinc pretreatment. We found no significant reduction in hair cell death in response to cotreatment alone (Figure 4A), and only a slight reduction at the highest zinc dose, 88 μg/L, in response to the one-hour pretreatment combined with cotreatment (Figure 4B). When treating fish with 88 μg/L zinc an hour before and in combination with a range of cadmium doses, we failed to see significant protection (Figure 4C). Zinc has also been shown to kill hair cells on its own (Montalbano et al., 2018), or to exacerbate hair cell death in response to other toxicants (Nakagawa et al., 1997); however, we saw no evidence of this at the doses we used (Figure 4).

**Figure 4:**
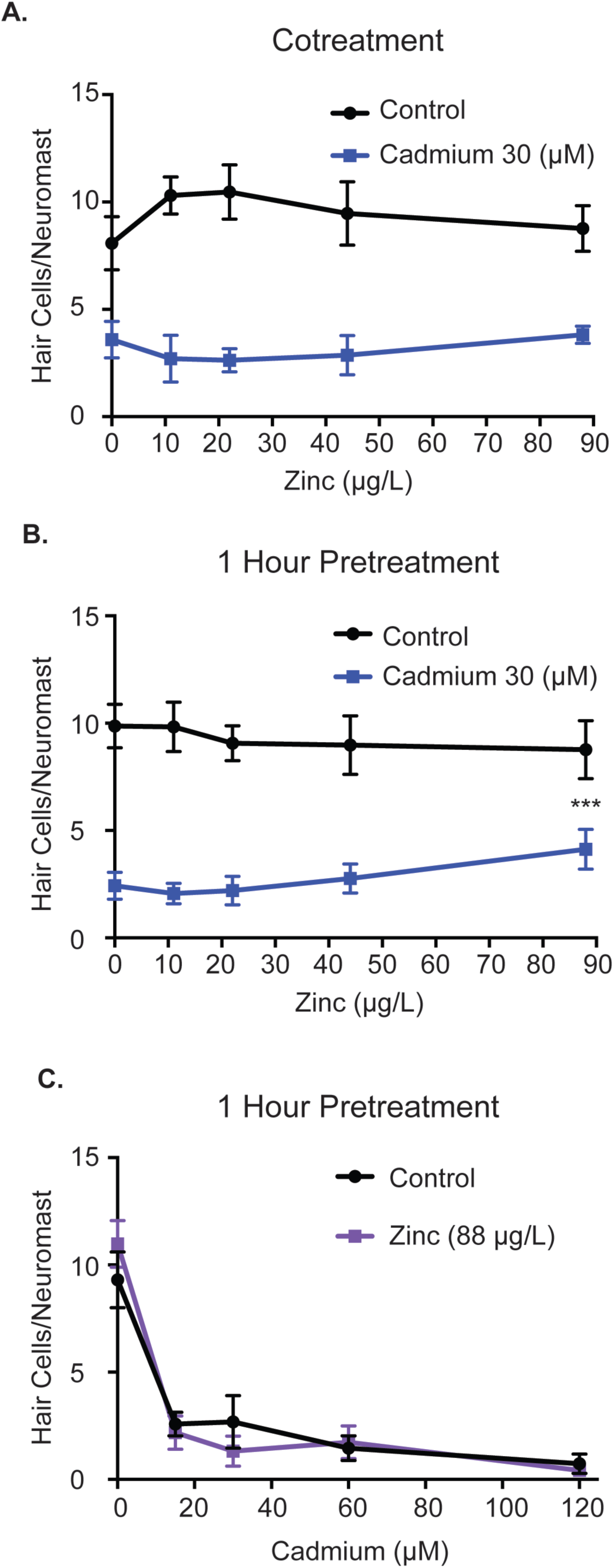
Zinc shows limited protection against cadmium-induced hair cell death. Zinc failed to reduce hair cell death when administered (A) alongside cadmium and only showed only a slight reduction at the highest dose tested when administered (B) one hour before and during cadmium treatment. This protection was not seen when tested against a range of cadmium doses (C). Data are displayed as mean ± standard deviation. *** = p<0.001 as compared to the 0 zinc control by Two-Way ANOVA and Šídák multiple comparisons test.

While cadmium is not believed to travel through copper transporter family gene products, it has been shown that copper cotreatment can impair cadmium uptake in fish (Komjarova and Bury, 2014). Copper has also been shown to protect against hair cell death in response to the platinum-based chemotherapeutic cisplatin and the heavy metal lead in mammals (Liu et al., 2011; More et al., 2010), though it was unable to protect against cisplatin in fish (Thomas et al., 2013). To determine whether copper might protect hair cells in zebrafish from cadmium toxicity, we first tested various doses of copper to see if they could protect against 30 μM of cadmium when fish were treated with copper for an hour before and while treated with cadmium. Copper has previously been shown to be toxic to zebrafish hair cells on its own (Hernández et al., 2006; Linbo et al., 2006; Mackenzie et al., 2012; Olivari et al., 2008) and we likewise found all but the smallest dose of copper tested, 0.25 μM, caused significant hair cell death on its own while offering no protection against cadmium (Figure 5A). We subsequently tested whether pre- and cotreatment with 0.25 μM copper would protect against a range of cadmium doses but again failed to see any significant protection (Figure 5B).

**Figure 5:**
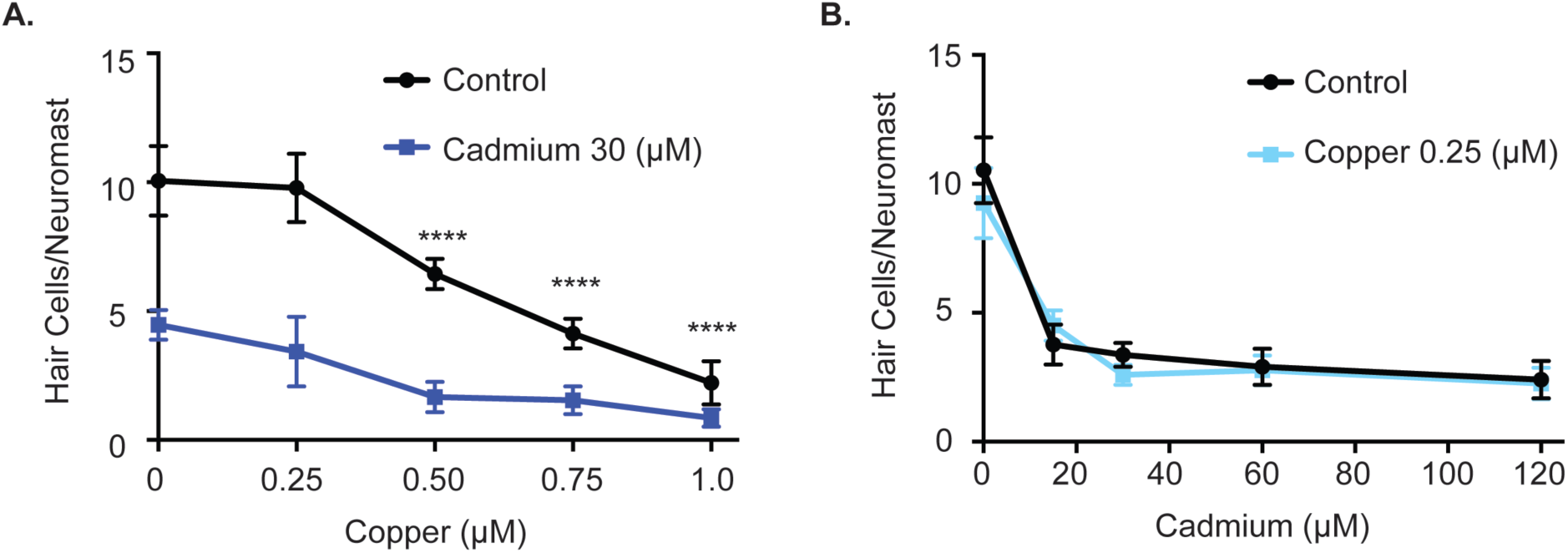
Copper fails to protect against cadmium-induced hair cell death. (A) At doses of 0.5 μM and higher copper caused significant hair cell death on its own. **** = p<0.0001 by Two-Way ANOVA and Šídák multiple comparisons test as compared to the 0-copper control. No dose of copper was able to protect against hair cell death in response to 30 μM of cadmium. (B) 0.25 μM of copper failed to show a significant reduction in hair cell death at all cadmium doses tested. Data are displayed as mean ± standard deviation.

## Discussion

Here we show that cadmium can consistently cause hair cell death in the lateral line system of zebrafish larvae. This is consistent with prior work that has shown cadmium can cause hair cell death in fish (Faucher et al., 2006; Montalbano et al., 2018; Sonnack et al., 2015). Studies looking at hair cell death and/or hearing loss in response to cadmium exposure in mammals and humans, on the other hand, have had much less consistent results, with some studies showing that cadmium is damaging (Choi et al., 2012; Choi and Park, 2017; Kim et al., 2008; Liu et al., 2014; Min et al., 2012; Ozcaglar et al., 2001) and others showing no effect (Carlson et al., 2018; Kang et al., 2018; Klimpel et al., 2017; Liu et al., 2018; Shiue, 2013; Whitworth et al., 1999). This disparity might come from the fact that lateral line hair cells in fish are on the surface of the animal and therefore easily accessible to toxins. In contrast to this, for cadmium to reach hair cells in mammals it must first travel through the bloodstream where it is actively removed by the liver and kidneys (Satarug, 2018; Swiergosz-Kowalewska, 2001; Yang and Shu, 2015), and then must pass through the blood-labyrinth barrier. This means presumably only a small percentage of the cadmium an individual is exposed to will ever make it to the hair cells. The fact that some researchers did see either hair cell damage or hearing impairment in mammals following cadmium exposure suggests that it can cause damage in these systems as long as sufficient amounts are present. Therefore, understanding how this damage occurs is an important question to research further.

In this study, we attempted to elucidate how cadmium enters hair cells. We did this by blocking potential uptake mechanisms and looking for a decrease in cadmium-induced hair cell toxicity. We showed that blocking mechanotransduction activity either via a genetic mutation in *cdh23* mutants or pharmacologically via benzamil lead to a significant reduction in cadmium-induced hair cell death. While *cdh23* mutants still showed significant cadmium-induced hair cell death at higher doses of cadmium, benzamil appeared to show an almost complete elimination of hair cell death across the dose range tested. This is different from what was previously shown for copper, where amiloride could protect against only lower doses of copper (Olivari et al., 2008). The reason for the differences in resistance seen in *cdh23* mutants versus benzamil treatment may be that unlike *cdh23*, benzamil is likely not blocking mechanotransduction selectively. Amiloride and benzamil are both considered nonselective epithelial sodium channel blockers and have been shown to be capable of blocking a number of channel types including some Ca^2+^, TRP, and K^+^ channels (Bielefeld et al., 1986; Castañeda et al., 2019; Dai et al., 2007; Tang et al., 1988). Therefore, while the mechanotransduction channel may serve as a major entry route for cadmium, cadmium could also be entering at a lower level through other channel types blocked by benzamil, thus allowing for the low level of hair cell death seen in cdh23 mutants. The small remaining level of hair cell death seen in *cdh23* mutants in the absence of mechanotransduction activity is in contrast to other toxicants entering via the mechanotransduction channel, such as cisplatin and aminoglycosides, where toxicity is completely blocked (Thomas et al., 2013; Wang and Steyger, 2009).

To test other potential uptake routes for cadmium into hair cells, we next investigated whether zinc or copper could protect hair cells from cadmium-induced hair cell death potentially be competing for entry. We failed to see consistent protection from either of these ions; however, these experiments were complicated by the fact that both copper and zinc can kill hair cells on their own (Hernández et al., 2006; Linbo et al., 2006; Montalbano et al., 2018), requiring the use of lower concentrations of these ions. Copper, in particular, kills hair cells at doses considerably lower than cadmium. The zinc doses we used were based on previous work showing that 22 μg/L zinc could protect against cadmium-induced olfactory damage whereas lower and higher doses could not (Heffern et al., 2018). We failed to see protection at 22 μg/L of zinc. We did see protection at the higher dose 88 μg/L zinc; however, that protection was inconsistent. Also, while benzamil was able to protect following cotreatment with cadmium, zinc required a one-hour pretreatment to show any effect.

While we failed to show consistent protection against cadmium-induced hair cell death from zinc, it has previously been shown that zinc can protect against cadmium-induced hearing loss in rats (Agirdir et al., 2002). Zinc can also protect against cadmium-induced nephrotoxicity (Liu et al., 1992; Tang et al., 1998). These experiments were looking over a longer period of cadmium treatment than we did. Potential mechanisms for this protection include the upregulation of metallothionein, changes in gene expression changes caused by cadmium that normally lead to toxicity, and prevention of cadmium from displacing zinc off metal-binding enzymes (Andrews, 2000; Pan et al., 2017; Pinter and Stillman, 2015). It is possible that these mechanisms do not have time to activate in the three-hour treatment window we are using. Also, we are exposing the fish to cadmium by adding it to the water they are swimming in. This is in contrast to mammalian studies that use more systemic treatments where cadmium is injected or given through drinking water. Presumably, given our short treatment times, cadmium is primarily exposed to the apical part of the hair cells where they are most accessible to the surrounding water. This may be the cause of the dominant role of mechanotransduction as a means of protecting against cadmium-induced hair cell death, as fewer other channels may be present on the apical area of the cell.

We also investigated whether two cilia-associated gene mutants that are resistant to aminoglycoside-induced hair cell death were similarly resistant to cadmium-induced hair cell death. Mutations in one of those genes, *ift88*, did lead to resistance in cadmium-induced hair cell death similar to what had previously been seen with aminoglycosides. These mutants have previously been shown to have a decrease in FM1-43 uptake (Stawicki et al., 2016) and a decreased response to water jet stimulation (Kindt et al., 2012), suggesting impaired mechanotransduction activity though mechanotransduction is not eliminated as in *cdh23* mutants where no response is seen (Nicolson et al., 1998). Thus, *ift88* mutant’s resistance to cadmium-induced hair cell death further supports the role of mechanotransduction in this process. They are not as resistant as *cdh23* mutants, which fits the fact that mechanotransduction is not as impaired. We failed to see significant resistance to cadmium-induced hair cell death in mutants of the other gene tested, *cc2d2a.* Unlike *ift88* mutants, these mutants do not show any impairment in FM1-43 uptake, suggesting normal mechanotransduction activity (Owens et al., 2008; Stawicki et al., 2016). They also do not show any impairments in aminoglycoside loading into hair cells (Owens et al., 2008; Stawicki et al., 2016), suggesting *cc2d2a* plays an intracellular role in the aminoglycoside-induced hair cell death process. The fact that these mutants are not resistant to cadmium-induced hair cell death suggests that these two toxins are behaving differently once entering the cell.

## Acknowledgements

We thank Amber-Lynn Lachowicz for zebrafish care.

## References

Agirdir, B. V., Bilgen, I., Dinc, O., Ozçaglar, H. Ü., Fisenk, F., Turhan, M., et al. (2002). Effect of Zinc Ion on Cadmium-Induced Auditory Changes. Biol. Trace Elem. Res. 88, 153–164. doi:10.1385/BTER:88:2:153.

Alharazneh, A., Luk, L., Huth, M., Monfared, A., Steyger, P. S., Cheng, A. G., et al. (2011). Functional hair cell mechanotransducer channels are required for aminoglycoside ototoxicity. PLoS One 6, e22347. doi:10.1371/journal.pone.0022347.

Andrews, G. K. (2000). Regulation of metallothionein gene expression by oxidative stress and metal ions. Biochem. Pharmacol. 59, 95–104. doi:10.1016/s0006-2952(99)00301-9.

Baker, C. F., and Montgomery, J. C. (2001). Sensory deficits induced by cadmium in banded kokopu, Galaxias fasciatus, juveniles. Environ. Biol. Fishes 62, 455–464. doi:10.1023/A:1012290912326.

Bannon, D. I., Abounader, R., Lees, P. S. J., and Bressler, J. P. (2003). Effect of DMT1 knockdown on iron, cadmium, and lead uptake in Caco-2 cells. Am. J. Physiol. Physiol. 284, C44–C50. doi:10.1152/ajpcell.00184.2002.

Barbier, O., Jacquillet, G., Tauc, M., Poujeol, P., and Cougnon, M. (2004). Acute study of interaction among cadmium, calcium, and zinc transport along the rat nephron in vivo. Am. J. Physiol. Physiol. 287, F1067–F1075. doi:10.1152/ajprenal.00120.2004.

Bielefeld, D. R., Hadley, R. W., Vassilev, P. M., and Hume, J. R. (1986). Membrane electrical properties of vesicular Na-Ca exchange inhibitors in single atrial myocytes. Circ. Res. 59, 381–9. doi:10.1161/01.res.59.4.381.

Brand, M., Heisenberg, C. P., Warga, R. M., Pelegri, F., Karlstrom, R. O., Beuchle, D., et al. (1996). Mutations affecting development of the midline and general body shape during zebrafish embryogenesis. Development 123, 129–142.

Carlson, K., Schacht, J., and Neitzel, R. L. (2018). Assessing ototoxicity due to chronic lead and cadmium intake with and without noise exposure in the mature mouse. J. Toxicol. Environ. Heal. Part A 81, 1041–1057. doi:10.1080/15287394.2018.1521320.

Castañeda, M. S., Tonini, R., Richards, C. D., Stocker, M., and Pedarzani, P. (2019). Benzamil inhibits neuronal and heterologously expressed small conductance Ca2+-activated K+ channels. Neuropharmacology 158, 107738. doi:10.1016/j.neuropharm.2019.107738.

Cheng, A. G., Cunningham, L. L., and Rubel, E. W. (2005). Mechanisms of hair cell death and protection. Curr. Opin. Otolaryngol. Head Neck Surg. 13, 343–8.

Choi, Y.-H., Hu, H., Mukherjee, B., Miller, J., and Park, S. K. (2012). Environmental cadmium and lead exposures and hearing loss in U.S. adults: the National Health and Nutrition Examination Survey, 1999 to 2004. Environ. Health Perspect. 120, 1544–50. doi:10.1289/ehp.1104863.

Choi, Y.-H., and Kim, K. (2014). Noise-Induced Hearing Loss in Korean Workers: Co-Exposure to Organic Solvents and Heavy Metals in Nationwide Industries. PLoS One 9, e97538. doi:10.1371/journal.pone.0097538.

Choi, Y.-H., and Park, S. K. (2017). Environmental Exposures to Lead, Mercury, and Cadmium and Hearing Loss in Adults and Adolescents: KNHANES 2010-2012. Environ. Health Perspect. 125, 067003. doi:10.1289/EHP565.

Dai, X.-Q., Ramji, A., Liu, Y., Li, Q., Karpinski, E., and Chen, X.-Z. (2007). Inhibition of TRPP3 Channel by Amiloride and Analogs. Mol. Pharmacol. 72, 1576–1585. doi:10.1124/mol.107.037150.

Dalton, T. P., He, L., Wang, B., Miller, M. L., Jin, L., Stringer, K. F., et al. (2005). Identification of mouse SLC39A8 as the transporter responsible for cadmium-induced toxicity in the testis. Proc. Natl. Acad. Sci. 102, 3401–3406. doi:10.1073/pnas.0406085102.

Faroon, O., Ashizawa, A., Wright, S., Tucker, P., Jenkins, K., Ingerman, L., et al. (2012). Toxicological Profile for Cadmium. Agency for Toxic Substances and Disease Registry (US), Atlanta (GA).

Faucher, K., Fichet, D., Miramand, P., and Lagardère, J.-P. (2008). Impact of chronic cadmium exposure at environmental dose on escape behaviour in sea bass (Dicentrarchus labrax L.; Teleostei, Moronidae). Environ. Pollut. 151, 148–157. doi:10.1016/J.ENVPOL.2007.02.017.

Faucher, K., Fichet, D., Miramand, P., and Lagardère, J. P. (2006). Impact of acute cadmium exposure on the trunk lateral line neuromasts and consequences on the “C-start” response behaviour of the sea bass (Dicentrarchus labrax L.; Teleostei, Moronidae). Aquat. Toxicol. 76, 278–294. doi:10.1016/J.AQUATOX.2005.10.004.

Ferraro, P. M., Costanzi, S., Naticchia, A., Sturniolo, A., and Gambaro, G. (2010). Low level exposure to cadmium increases the risk of chronic kidney disease: analysis of the NHANES 1999-2006. BMC Public Health 10, 304. doi:10.1186/1471-2458-10-304.

Fujishiro, H., Okugaki, S., Kubota, K., Fujiyama, T., Miyataka, H., and Himeno, S. (2009). The role of ZIP8 down-regulation in cadmium-resistant metallothionein-null cells. J. Appl. Toxicol. 29, 367–373. doi:10.1002/jat.1419.

Gallagher, C. M., Kovach, J. S., and Meliker, J. R. (2008). Urinary Cadmium and Osteoporosis in U.S. Women ≥50 Years of Age: NHANES 1988–1994 and 1999– 2004. Environ. Health Perspect. 116, 1338–1343. doi:10.1289/ehp.11452.

García-Esquinas, E., Pollan, M., Tellez-Plaza, M., Francesconi, K. A., Goessler, W., Guallar, E., et al. (2014). Cadmium Exposure and Cancer Mortality in a Prospective Cohort: The Strong Heart Study. Environ. Health Perspect. 122, 363–370. doi:10.1289/ehp.1306587.

Girijashanker, K., He, L., Soleimani, M., Reed, J. M., Li, H., Liu, Z., et al. (2008). Slc39a14 Gene Encodes ZIP14, A Metal/Bicarbonate Symporter: Similarities to the ZIP8 Transporter. Mol. Pharmacol. 73, 1413–1423. doi:10.1124/mol.107.043588.

Goman, A. M., and Lin, F. R. (2016). Prevalence of Hearing Loss by Severity in the United States. Am. J. Public Health 106, 1820–1822. doi:10.2105/AJPH.2016.303299.

Hailey, D. W., Esterberg, R., Linbo, T. H., Rubel, E. W., and Raible, D. W. (2017). Fluorescent aminoglycosides reveal intracellular trafficking routes in mechanosensory hair cells. J. Clin. Invest. 127, 472–486. doi:10.1172/JCI85052.

Harris, J. A., Cheng, A. G., Cunningham, L. L., MacDonald, G., Raible, D. W., and Rubel, E. W. (2003). Neomycin-induced hair cell death and rapid regeneration in the lateral line of zebrafish (Danio rerio). J. Assoc. Res. Otolaryngol. 4, 219–234. doi:10.1007/s10162-002-3022-x.

Heffern, K., Tierney, K., and Gallagher, E. P. (2018). Comparative effects of cadmium, zinc, arsenic and chromium on olfactory-mediated neurobehavior and gene expression in larval zebrafish (Danio rerio). Aquat. Toxicol. 201, 83–90. doi:10.1016/j.aquatox.2018.05.016.

Hernández, P. P., Moreno, V., Olivari, F. A., and Allende, M. L. (2006). Sub-lethal concentrations of waterborne copper are toxic to lateral line neuromasts in zebrafish (Danio rerio). Hear. Res. 213, 1–10. doi:10.1016/j.heares.2005.10.015.

Hyder, O., Chung, M., Cosgrove, D., Herman, J. M., Li, Z., Firoozmand, A., et al. (2013). Cadmium Exposure and Liver Disease among US Adults. J. Gastrointest. Surg. 17, 1265–1273. doi:10.1007/s11605-013-2210-9.

IARC (2012). Cadmium and cadmium compounds. IARC 100C, 121–145. doi:10.1002/14356007.a04.

Kang, G. H., Uhm, J. Y., Choi, Y. G., Kang, E. K., Kim, S. Y., Choo, W. O., et al. (2018). Environmental exposure of heavy metal (lead and cadmium) and hearing loss: data from the Korea National Health and Nutrition Examination Survey (KNHANES 2010–2013). Ann. Occup. Environ. Med. 30, 22. doi:10.1186/s40557-018-0237-9.

Kim, S.-J., Jeong, H.-J., Myung, N.-Y., Kim, M., Lee, J.-H., So, H., et al. (2008). The protective mechanism of antioxidants in cadmium-induced ototoxicity in vitro and in vivo. Environ. Health Perspect. 116, 854–62. doi:10.1289/ehp.10467.

Kindt, K. S., Finch, G., and Nicolson, T. (2012). Kinocilia mediate mechanosensitivity in developing zebrafish hair cells. Dev. Cell 23, 329–341. doi:10.1016/j.devcel.2012.05.022.

Klimpel, K. E. M., Lee, M. Y., King, W. M., Raphael, Y., Schacht, J., and Neitzel, R. L. (2017). Vestibular dysfunction in the adult CBA/CaJ mouse after lead and cadmium treatment. Environ. Toxicol. 32, 869–876. doi:10.1002/tox.22286.

Komjarova, I., and Bury, N. R. (2014). Evidence of Common Cadmium and Copper Uptake Routes in Zebrafish *Danio rerio*. Environ. Sci. Technol. 48, 12946–12951. doi:10.1021/es5032272.

Kovacs, G., Danko, T., Bergeron, M. J., Balazs, B., Suzuki, Y., Zsembery, A., et al. (2011). Heavy metal cations permeate the TRPV6 epithelial cation channel. Cell Calcium 49, 43–55. doi:10.1016/j.ceca.2010.11.007.

Kovacs, G., Montalbetti, N., Franz, M.-C., Graeter, S., Simonin, A., and Hediger, M. A. (2013). Human TRPV5 and TRPV6: Key players in cadmium and zinc toxicity. Cell Calcium 54, 276–286. doi:10.1016/j.ceca.2013.07.003.

Kurabi, A., Keithley, E. M., Housley, G. D., Ryan, A. F., and Wong, A. C.-Y. (2017). Cellular mechanisms of noise-induced hearing loss. Hear. Res. 349, 129–137. doi:10.1016/j.heares.2016.11.013.

Larsson, M. (2012). Why do fish school? Curr. Zool. 58, 116–128. doi:10.1093/czoolo/58.1.116.

Lévesque, M., Martineau, C., Jumarie, C., and Moreau, R. (2008). Characterization of cadmium uptake and cytotoxicity in human osteoblast-like MG-63 cells. Toxicol. Appl. Pharmacol. 231, 308–317. doi:10.1016/j.taap.2008.04.016.

Lin, F. R., Niparko, J. K., and Ferrucci, L. (2011). Hearing loss prevalence in the United States. Arch. Intern. Med. 171, 1851–1852. doi:10.1001/archinternmed.2011.506.

Linbo, T. L., Stehr, C. M., Incardona, J. P., and Scholz, N. L. (2006). Dissolved copper triggers cell death in the peripheral mechanosensory system of larval fish. Environ. Toxicol. Chem. 25, 597–603.

Liu, H., Ding, D., Sun, H., Jiang, H., Wu, X., Roth, J. A., et al. (2014). Cadmium-induced ototoxicity in rat cochlear organotypic cultures. Neurotox. Res. 26, 179–89. doi:10.1007/s12640-014-9461-4.

Liu, S., Zhang, K., Wu, S., Ji, X., Li, N., Liu, R., et al. (2011). Lead-Induced Hearing Loss in Rats and the Protective Effect of Copper. Biol. Trace Elem. Res. 144, 1112–1119. doi:10.1007/s12011-011-9142-6.

Liu, X. Y., Jin, T. Y., Nordberg, G. F., Rännar, S., Sjöström, M., and Zhou, Y. (1992). A multivariate study of protective effects of Zn and Cu against nephrotoxicity induced by cadmium metallothionein in rats. Toxicol. Appl. Pharmacol. 114, 239–45. doi:10.1016/0041-008x(92)90074-3.

Liu, Y., Huo, X., Xu, L., Wei, X., Wu, W., Wu, X., et al. (2018). Hearing loss in children with e-waste lead and cadmium exposure. Sci. Total Environ. 624, 621–627. doi:10.1016/j.scitotenv.2017.12.091.

Low, J., and Higgs, D. M. (2015). Sublethal effects of cadmium on auditory structure and function in fathead minnows (Pimephales promelas). Fish Physiol. Biochem. 41, 357–369. doi:10.1007/s10695-014-9988-6.

Mackenzie, S. M., Raible, D. W., Nguyen, T., Hume, C., Oesterle, E., and Raible, D. (2012). Proliferative Regeneration of Zebrafish Lateral Line Hair Cells after Different Ototoxic Insults. PLoS One 7, e47257. doi:10.1371/journal.pone.0047257.

Marcotti, W., van Netten, S. M., and Kros, C. J. (2005). The aminoglycoside antibiotic dihydrostreptomycin rapidly enters mouse outer hair cells through the mechano-electrical transducer channels. J. Physiol. 567, 505–521. doi:10.1113/jphysiol.2005.085951.

Martineau, C., Abed, E., Médina, G., Jomphe, L.-A., Mantha, M., Jumarie, C., et al. (2010). Involvement of transient receptor potential melastatin-related 7 (TRPM7) channels in cadmium uptake and cytotoxicity in MC3T3-E1 osteoblasts. Toxicol. Lett. 199, 357–363. doi:10.1016/j.toxlet.2010.09.019.

Min, K.-B., Lee, K.-J., Park, J.-B., and Min, J.-Y. (2012). Lead and Cadmium Levels and Balance and Vestibular Dysfunction among Adult Participants in the National Health and Nutrition Examination Survey (NHANES) 1999–2004. Environ. Health Perspect. 120, 413–417. doi:10.1289/ehp.1103643.

Montalbano, G., Capillo, G., Laurà, R., Abbate, F., Levanti, M., Guerrera, M. C., et al. (2018). Neuromast hair cells retain the capacity of regeneration during heavy metal exposure. Ann. Anat. - Anat. Anzeiger 218, 183–189. doi:10.1016/j.aanat.2018.03.007.

More, S. S., Akil, O., Ianculescu, A. G., Geier, E. G., Lustig, L. R., and Giacomini, K. M. (2010). Role of the Copper Transporter, CTR1, in Platinum-Induced Ototoxicity. J. Neurosci. 30, 9500–9509. doi:10.1523/JNEUROSCI.1544-10.2010.

Nakagawa, T., Yamane, H., Shibata, S., Sunami, K., and Nakai, Y. (1997). Cell death caused by the acute effects of aminoglycoside and zinc in the ampullary cristae of guinea pigs. Eur. Arch. Otorhinolaryngol. 254, 153–7.

Nicolson, T., Rüsch, A., Friedrich, R. W., Granato, M., Ruppersberg, J. P., and Nüsslein-Volhard, C. (1998). Genetic Analysis of Vertebrate Sensory Hair Cell Mechanosensation: the Zebrafish Circler Mutants. Neuron 20, 271–283. doi:10.1016/S0896-6273(00)80455-9.

Olivari, F. A., Hernández, P. P., and Allende, M. L. (2008). Acute copper exposure induces oxidative stress and cell death in lateral line hair cells of zebrafish larvae. Brain Res. 1244, 1–12. doi:10.1016/J.BRAINRES.2008.09.050.

Olivi, L., Sisk, J., and Bressler, J. (2001). Involvement of DMT1 in uptake of Cd in MDCK cells: role of protein kinase C. Am. J. Physiol. Physiol. 281, C793–C800. doi:10.1152/ajpcell.2001.281.3.C793.

Ou, H. C., Raible, D. W., and Rubel, E. W. (2007). Cisplatin-induced hair cell loss in zebrafish (Danio rerio) lateral line. Hear. Res. 233, 46–53. doi:10.1016/j.heares.2007.07.003.

Owens, K. N., Santos, F., Roberts, B., Linbo, T., Coffin, A. B., Knisely, A. J., et al. (2008). Identification of genetic and chemical modulators of zebrafish mechanosensory hair cell death. PLoS Genet. 4, e1000020. doi:10.1371/journal.pgen.1000020.

Ozcaglar, H. U., Agirdir, B., Dinc, O., Turhan, M., Kilinçarslan, S., and Oner, G. (2001). Effects of cadmium on the hearing system. Acta Otolaryngol. 121, 393–7.

Pan, J., Huang, X., Li, Y., Li, M., Yao, N., Zhou, Z., et al. (2017). Zinc protects against cadmium-induced toxicity by regulating oxidative stress, ions homeostasis and protein synthesis. Chemosphere 188, 265–273. doi:10.1016/j.chemosphere.2017.08.106.

Pinter, T. B. J., and Stillman, M. J. (2015). Kinetics of Zinc and Cadmium Exchanges between Metallothionein and Carbonic Anhydrase. Biochemistry 54, 6284–6293. doi:10.1021/acs.biochem.5b00912.

Raible, D. W., and Kruse, G. J. (2000). Organization of the lateral line system in embryonic zebrafish. J. Comp. Neurol. 421, 189–198.

Satarug, S. (2018). Dietary Cadmium Intake and Its Effects on Kidneys. Toxics 6. doi:10.3390/toxics6010015.

Schaal, N., Slagley, J., Zreiqat, M., and Paschold, H. (2017). Effects of combined exposure to metals, solvents, and noise on permanent threshold shifts. Am. J. Ind. Med. 60, 227–238. doi:10.1002/ajim.22690.

Schacht, J., Talaska, A. E., and Rybak, L. P. (2012). Cisplatin and Aminoglycoside Antibiotics: Hearing Loss and Its Prevention. Anat. Rec. Adv. Integr. Anat. Evol. Biol. 295, 1837–1850. doi:10.1002/ar.22578.

Seiler, C., and Nicolson, T. (1999). Defective calmodulin-dependent rapid apical endocytosis in zebrafish sensory hair cell mutants. J. Neurobiol. 41, 424–434.

Shiue, I. (2013). Urinary environmental chemical concentrations and vitamin D are associated with vision, hearing, and balance disorders in the elderly. Environ. Int. 53, 41–46. doi:10.1016/j.envint.2012.12.006.

Söllner, C., Rauch, G.-J., Siemens, J., Geisler, R., Schuster, S. C., Müller, U., et al. (2004). Mutations in cadherin 23 affect tip links in zebrafish sensory hair cells. Nature 428, 955–959. doi:10.1038/nature02484.

Sonnack, L., Kampe, S., Muth-Köhne, E., Erdinger, L., Henny, N., Hollert, H., et al. (2015). Effects of metal exposure on motor neuron development, neuromasts and the escape response of zebrafish embryos. Neurotoxicol. Teratol. 50, 33–42. doi:10.1016/j.ntt.2015.05.006.

Soodvilai, S., Nantavishit, J., Muanprasat, C., and Chatsudthipong, V. (2011). Renal organic cation transporters mediated cadmium-induced nephrotoxicity. Toxicol. Lett. 204, 38–42. doi:10.1016/j.toxlet.2011.04.005.

Stawicki, T. M., Hernandez, L., Esterberg, R., Linbo, T., Owens, K. N., Shah, A. N., et al. (2016). Cilia-associated genes play differing roles in aminoglycoside-induced hair cell death in zebrafish. G3 Genes, Genomes, Genet. 6, 2225–2235. doi:10.1534/g3.116.030080.

Stawicki, T. M., Owens, K. N., Linbo, T., Reinhart, K. E., Rubel, E. W., and Raible, D. W. (2014). The zebrafish merovingian mutant reveals a role for pH regulation in hair cell toxicity and function. Dis. Model. Mech. 7, 847–856. doi:10.1242/dmm.016576.

Swiergosz-Kowalewska, R. (2001). Cadmium distribution and toxicity in tissues of small rodents. Microsc. Res. Tech. 55, 208–222. doi:10.1002/jemt.1171.

Tang, C. M., Presser, F., and Morad, M. (1988). Amiloride selectively blocks the low threshold (T) calcium channel. Science 240, 213–5. doi:10.1126/science.2451291.

Tang, W., Sadovic, S., and Shaikh, Z. A. (1998). Nephrotoxicity of Cadmium-Metallothionein: Protection by Zinc and Role of Glutathione. Toxicol. Appl. Pharmacol. 151, 276–282. doi:10.1006/taap.1998.8465.

Tellez-Plaza, M., Navas-Acien, A., Menke, A., Crainiceanu, C. M., Pastor-Barriuso, R., and Guallar, E. (2012). Cadmium Exposure and All-Cause and Cardiovascular Mortality in the U.S. General Population. Environ. Health Perspect. 120, 1017–1022. doi:10.1289/ehp.1104352.

Thomas, A. J., Hailey, D. W., Stawicki, T. M., Wu, P., Coffin, A. B., Rubel, E. W., et al. (2013). Functional mechanotransduction is required for cisplatin-induced hair cell death in the zebrafish lateral line. J. Neurosci. 33, 4405–14. doi:10.1523/JNEUROSCI.3940-12.2013.

Ton, C., and Parng, C. (2005). The use of zebrafish for assessing ototoxic and otoprotective agents. Hear. Res. 208, 79–88. doi:10.1016/j.heares.2005.05.005.

Tsujikawa, M., and Malicki, J. (2004). Intraflagellar Transport Genes Are Essential for Differentiation and Survival of Vertebrate Sensory Neurons. Neuron 42, 703–716. doi:10.1016/S0896-6273(04)00268-5.

Wang, L., and Gallagher, E. P. (2013). Role of Nrf2 antioxidant defense in mitigating cadmium-induced oxidative stress in the olfactory system of zebrafish. Toxicol. Appl. Pharmacol. 266, 177–186. doi:10.1016/j.taap.2012.11.010.

Wang, Q., and Steyger, P. S. (2009). Trafficking of Systemic Fluorescent Gentamicin into the Cochlea and Hair Cells. J. Assoc. Res. Otolaryngol. 10, 205–219. doi:10.1007/s10162-009-0160-4.

Whitworth, C. A., Hudson, T. E., and Rybak, L. P. (1999). The effect of combined administration of cadmium and furosemide on auditory function in the rat. Hear. Res. 129, 61–70. doi:10.1016/S0378-5955(98)00222-6.

Yamasoba, T., Lin, F. R., Someya, S., Kashio, A., Sakamoto, T., and Kondo, K. (2013). Current concepts in age-related hearing loss: Epidemiology and mechanistic pathways. Hear. Res. 303, 30–38. doi:10.1016/j.heares.2013.01.021.

Yang, H., and Shu, Y. (2015). Cadmium transporters in the kidney and cadmium-induced nephrotoxicity. Int. J. Mol. Sci. 16, 1484–1494. doi:10.3390/ijms16011484.

